# CDK4/6 inhibitors induce replication stress to cause long-term cell cycle withdrawal

**DOI:** 10.1101/2021.02.03.428245

**Authors:** Lisa Crozier, Reece Foy, Brandon L. Mouery, Robert H. Whitaker, Andrea Corno, Christos Spanos, Tony Ly, Jeanette Gowen Cook, Adrian T. Saurin

**Author notes:** These authors contributed equally (Lisa Crozier, Reece Foy).

## Abstract

CDK4/6 inhibitors arrest the cell cycle in G1-phase. They are approved to treat breast cancer and are also undergoing clinical trials against a range of other tumour types. To facilitate these efforts, it is important to understand why a cytostatic arrest in G1 causes long-lasting effects on tumour growth. Here we demonstrate that a prolonged G1-arrest following CDK4/6 inhibition downregulates replisome components and impairs origin licencing. This causes a failure in DNA replication after release from that arrest, resulting in a p53-dependent withdrawal from the cell cycle. If p53 is absent, then cells bypass the G2-checkpoint and undergo a catastrophic mitosis resulting in excessive DNA damage. These data therefore link CDK4/6 inhibition to genotoxic stress; a phenotype that is shared by most other broad-spectrum anti-cancer drugs. This provides a rationale to predict responsive tumour types and effective combination therapies, as demonstrated by the fact that CDK4/6 inhibition induces sensitivity to chemotherapeutics that also cause replication stress.

## Introduction

Cyclin-dependent kinases 4 and 6 (CDK4/6) phosphorylate the retinoblastoma protein (Rb) to relieve repression of E2F-dependent genes and allow progression from G1 into S-phase. Three structurally distinct CDK4/6 inhibitors have recently been licenced for breast cancer treatment: palbociclib, ribociclib and abemaciclib ^1,2^. Unlike other cell cycle inhibitors, these agents are generally well-tolerated and have demonstrated remarkable efficacy in treating hormone receptor-positive (HR+), human epidermal growth factor 2-negative (HER2-) metastatic breast cancer ^3-6^. Comparisons with standard of care chemotherapy have given weight to the notion that CDK4/6 inhibitors may be able to replace conventional chemotherapy in this cancer subtype, which represents the majority of metastatic breast cancers ^7-9^.

There is also a wealth of preclinical evidence that CDK4/6 inhibitors display broad activity against a wide range of other tumour types (for reviews see ^10-12^). This is supported by preliminary clinical data suggesting that these inhibitors may be beneficial for treating non-small cell lung cancer, melanoma, head and neck squamous cell carcinoma, mantle cell lymphoma, triple-negative breast cancer and acute myeloid leukaemia ^10,13-17^. Currently, there are at least 18 different CDK4/6 inhibitors being tested in over 100 clinical trials against various tumour types (for reviews see: ^2,11,12,18,19^). The hope is that these targeted cell cycle inhibitors may be widely applicable for cancer treatment, perhaps offering an alternative to the non-targeted, and considerably more toxic, DNA damaging agents or microtubule poisons that are currently in widespread clinical use.

To facilitate these efforts, there is an urgent need to identify biomarkers and combination treatments that can predict and improve patient outcomes. This requires the characterisation of sensitizing events that can either: 1) enhance the ability of CDK4/6 inhibitors to arrest the cell cycle in G1, or 2) improve long-term outcomes following this G1 arrest. Although various genetic backgrounds and drug treatments are known to sensitize the CyclinD-CDK4/6-Rb pathway and promote an efficient G1-arrest ^1,2,20-24^}, relatively little is known about sensitizing events that could improve long-term growth suppression following this arrest. The problem is that there is no clear consensus for why a G1 arrest, which is essentially cytostatic, can produce durable effects in patients. There are many different potential explanations, including that CDK4/6 inhibition can induce senescence, apoptosis, metabolic reprogramming and/or anti-tumour immunity (for reviews see: ^18,24^), but whether a common event underlies these different outcomes is unclear. There is good evidence that some of the long-term outcomes are linked, in particular, senescent cells can secrete a variety of factors that engage the immune system ^25-28^, and this senescence phenotype contributes to the ability of CDK4/6 inhibition to sensitize tumours to immune checkpoint blockade ^29-31^. It is therefore critical to determine how and why G1-arrested cells eventually become senescent because this may help to inform ongoing clinical trials assessing CDK4/6 inhibition alongside immunotherapy (currently 14 trials in 8 different cancer types ^32^).

Senescence is a state of irreversible cell cycle exit induced by stress, typically DNA damage or oxidative stress ^33^. A crucial question therefore concerns the nature of the stress that leads to senescence following CDK4/6 inhibition. Unfortunately, although senescence has been demonstrated in a variety of different studies (for recent review see: ^32^), only two of these studies report a source for the stress. In both cases, senescence is believed to be induced by ROS generated during a G1 arrest ^20^, perhaps as a result of FOXM1 destabilisation ^34^. There have been more attempts to characterise the mediator(s) of the subsequent senescent response, but the answers here have been varied, including a dependence on ATRX ^35,36^, proteasome activation ^37^, mTOR activation ^38^ or mTOR inhibition ^39^. This variability may reflect inherent differences between genomically diverse cancer lines. Alternatively, it may be due to inconsistent treatment protocols (drug type, dose, duration of exposure and length of washout) or the reliance on fixed endpoints that can only indirectly measure senescence ^40^.

To overcome these problems, we elected to use a non-transformed near-diploid RPE1 cell line expressing a FUCCI cell cycle reporter to track the fate of single cells over time following CDK4/6 inhibition. We compared all currently licenced CDK4/6 inhibitors over a range of treatment protocols to address one key unexplained question: why do these inhibitors cause long-term cell cycle exit? Our results demonstrate that a prolonged G1-arrest is associated with the downregulation of replisome components, including the MCM complex, which causes reduced origin licencing, replication stress, p53-p21 activation and long-term cell cycle withdrawal. These results therefore identify a major source of genotoxic stress induced by targeted CDK4/6 inhibition. This finding has considerable implications for the identification of sensitive/resistant tumour types, for the design of effective combination treatments and drug dosing schedules, and for the efforts to use CDK4/6 inhibitors to sensitize tumours to immune checkpoint blockade.

## Results

We first quantified the fraction of G1-arrested RPE1-FUCCI cells ^41^ following 24h treatment with four structurally distinct CDK4/6 inhibitors: palbociclib (PD-0332991), ribociclib (LEE-011), abemaciclib (LY-2835219) and trilaciclib (G1T28). The dose-response curves for all inhibitors demonstrate a penetrant arrest at the clinically-relevant peak plasma concentrations (Cmax) observed in patients ^18,42^ (Figure 1A). Note, RPE1 cells are exquisitely sensitive to these compounds since the IC50s for palbociclib and abemaciclib (150nM and 65nM, respectively) were comparable to the IC50 values reported in the most sensitive cell type from a panel of 560 tumour lines: 130nM palbociclib in MDA-MB-175-VII (breast cancer) and 60nM abemaciclib in JeKo-1 (mantle cell lymphoma) ^21^. At approximate Cmax concentrations or lower, the G1 arrest was fully reversible within 24h of drug washout; although at higher concentrations this reversibility is compromised for all drugs (Figure 1A). Note, that we used an extensive washout protocol to ensure that persistent arrest is due to effects on the cell cycle and not incomplete drug washout (Figure S1; this protocol was used in all subsequent washout experiments). The irreversible effects at higher drug concentrations are likely to represent off-target effects, as reported previously for palbociclib at ≥ 5μM concentrations ^20^. In general, abemaciclib displayed the narrowest concentration range in which to achieve an efficient arrest that remained reversible, as noted recent by others ^43^. The fact that abemaciclib is uniquely able to induce irreversible effects at approximate Cmax concentrations may help to explain the unique toxicity profile associated with this drug ^44^.

**Figure 1:**
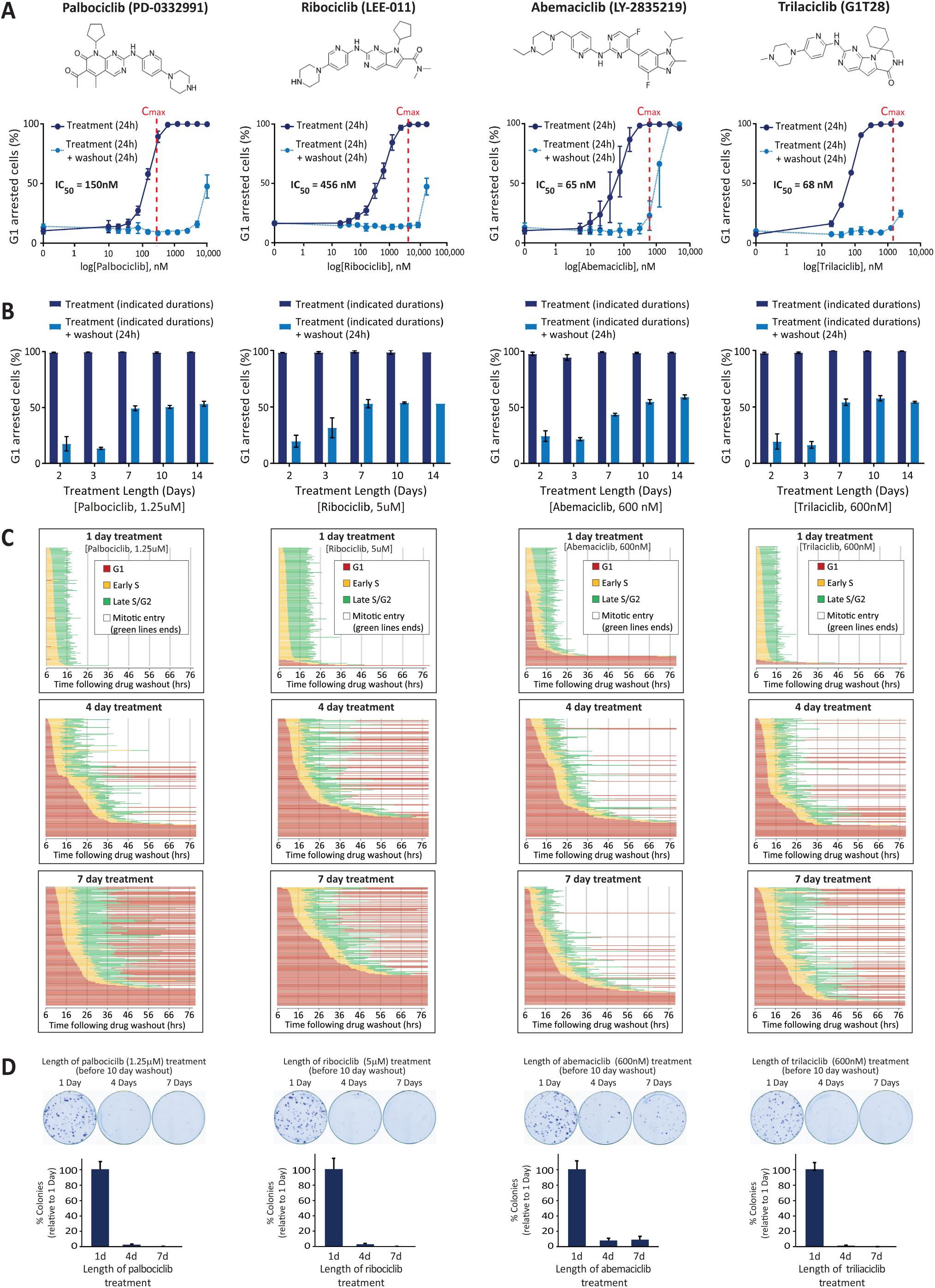
A prolonged G1 arrest following CDK4/6 inhibition causes defects in the next cell cycle. **(A)** Top panel displays structure of each CDK4/6 inhibitor tested. Bottom panel shows dose response curves with these inhibitors displaying percentage of G1-arrested RPE1-FUCCI cells. To obtain dose response curves, the number of red (G1-arrested) cells were calculated following 24hr drug addition (dark blue solid lines) or 24h after subsequent drug washout (light blue dotted lines). Cmax values observed in patients (taken from ^18,42^) are represented on each graph with red dotted lines. Graphs display mean data -/+ SEM from 3 experiments, with at least 500 cells counted per condition per experiment. **(B)** Percentage of G1-arrested RPE1-FUCCI cells, calculated as in panel A, but using a fixed concentration of CDK4/6 inhibitor for different durations of time, as indicated. Each bar displays mean data -/+ SEM from 3 experiments, with at least 500 cells counted per condition per experiment. **(C)** Cell cycle profile of individual RPE1-FUCCI cells (each bar represents one cell) after washout from 1 (top panel), 4 (middle panel) or 7 (bottom panel) days of treatment with CDK4/6 inhibitor, at the indicated doses (same concentration used in panel B). STLC (10 μM) was added to prevent progression past the first mitosis. 50 cells were analysed at random for each repeat and 3 experimental repeats are displayed (150 cells total). **(D)** Top panel shows representative images of colony forming assays of RPE1 cells treated with CDK4/6 inhibitor for 1, 4 or 7 days and then grown at low density without inhibitor for 10 days. Inhibitors were added at the same concentrations used in panel B. Bottom panel shows the quantification of these images. Each bar displays mean data -/+ SEM from 3 experiments.

Increasing the duration of drug exposure to 48h produced almost identical dose-response curves, indicating that after 2 days of treatment all drugs induced a similarly reversible G1 arrest (Figure S2). We next used the minimal dose of each drug required to produce a fully penetrant G1 arrest for 24h, and assessed the ability of this dose to induce a prolonged arrest for up to 14 days. Figure 1B demonstrates that all drugs can hold a full G1-arrest for up to 2 weeks. However, upon release from a prolonged arrest (> 3 days), we observed an increase in the fraction of cells remaining in G1. Therefore, CDK4/6 inhibition can induce a penetrant and reversible cell cycle arrest in RPE1 cells, but this reversibility is compromised when drug treatment persists for longer than 3 days.

To analyse this phenotype more closely, we performed live single-cell fate analysis using RPE1-FUCCI cells during the first cell cycle after washout from different durations of CDK4/6 inhibitor treatment (Figure 1C) ^45^. Using this approach, we observed two striking phenotypes that appeared specifically following release from prolonged drug exposure. Firstly, the length of time individual cells took to exit G1 and enter S-phase following drug washout increased: most cells took many hours to exit G1 and a small fraction of cells failed to exit G1 at all within the 3-day imaging period. This is suggestive of a deep G1 arrest, which may become irreversible in a subset of cells. Secondly, following washout from 4 and 7-day treatments, many cells that entered S-phase failed to reach mitosis and instead reverted back into a G1-like state: green bars turning red (G1) instead of white (mitosis) in Figure 1C. This was not due to depletion of nutrients in the media since it was unaffected by replenishing the media daily (Figure S3). Therefore, prolonged arrest with CDK4/6 inhibitors induces a deep G1-arrest, and many cells that exit from this arrest fail to complete the next cell cycle. Colony forming assay demonstrated that these effects are associated with long-term inhibition of cell growth (Figure 1D).

The reversion of cells from S-phase/G2 back into G1 has previously been associated with a p53-dependent senescent response ^41,46,47^. To explore the role of p53 in these phenotypes, we performed similar cell fate analysis in p53-WT and p53-KO RPE-FUCCI cells, generated by CRISPR/Cas9-mediated gene-editing (Figure S4). Figure 2A demonstrates that 24h palbociclib treatment induced a dose-dependent reversible G1-arrest that was indistinguishable between p53-WT and KO cells. Although knockout of p53 did not affect the efficiency of a palbociclib-induced arrest, it did produce a striking effect on the phenotypes observed following washout from prolonged drug exposure (Figure 2B). Firstly, the delay in S-phase entry following drug washout was less pronounced and fewer cells remained arrested in G1 for the duration of the movie. Secondly, the conversions from S-phase/G2 into G1 were completely abrogated.

**Figure 2.**
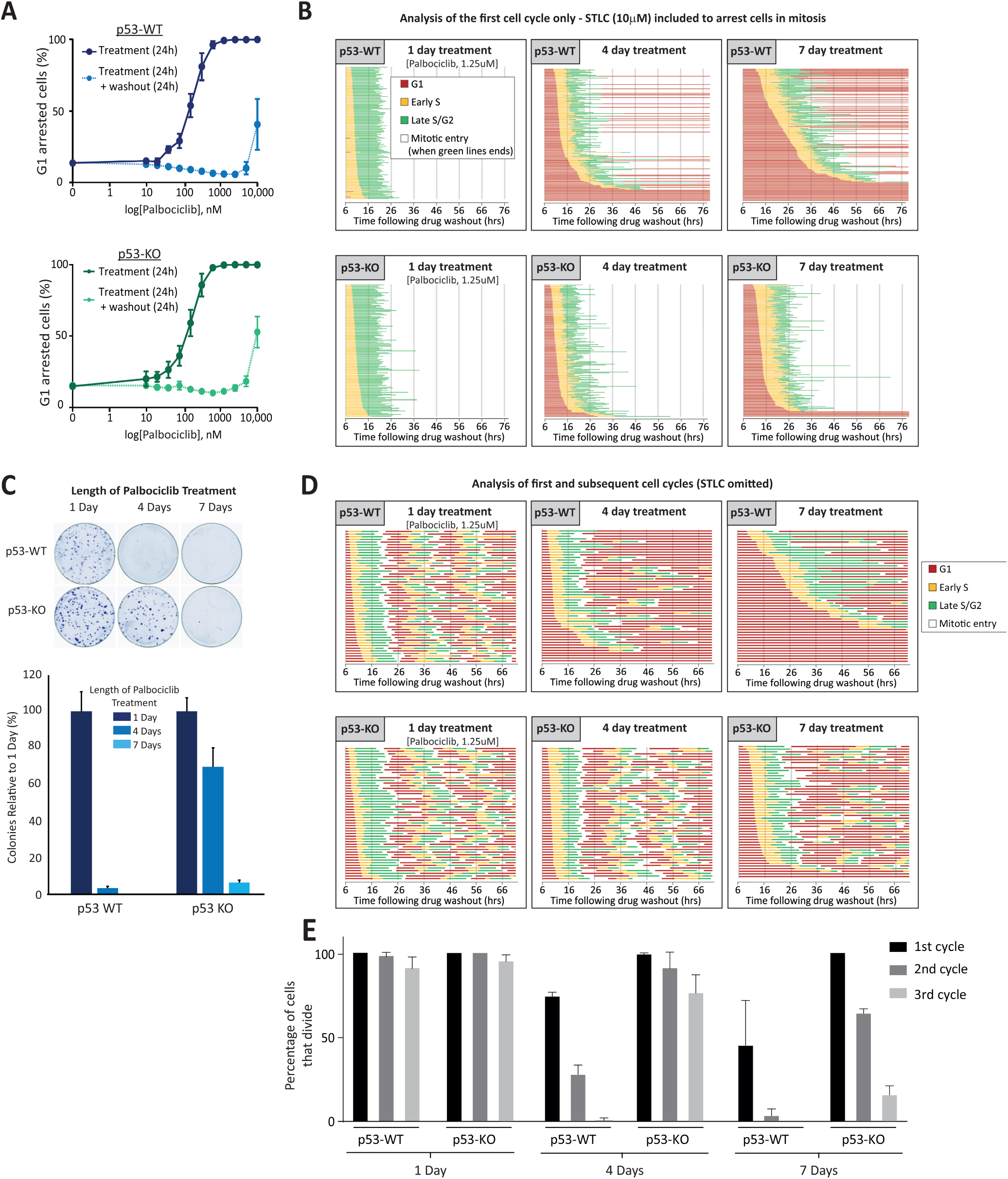
P53 loss restores cell cycle progression and enhances long-term growth following prolonged CDK4/6 inhibition. **(A)** Dose response curves displaying the percentage of G1-arrested p53-WT (blue) or KO (green) RPE1-FUCCI cells following 24hr incubation with palbociclib (dark solid lines) or 24h after subsequent washout (light dotted lines). Graphs display mean data -/+ SEM from 3 experiments, with at least 500 cells counted per condition per experiment. **(B)** Cell cycle profile of individual p53-WT or KO RPE1-FUCCI cells (each bar represents one cell) after washout from 1, 4 or 7 days of treatment with palbociclib (1.25μM). STLC (10 μM) was added to prevent progression past the first mitosis. 50 cells were analysed at random for each repeat and 3 experimental repeats are displayed (150 cells total). **(C)** Top panel shows representative images of colony forming assays in p53-WT or KO RPE1 cells treated with palbociclib (1.25μM) for 1, 4 or 7 days and then grown at low density without inhibitor for 10 days. Bottom panel shows the quantification of these images. Each bar displays mean data -/+ SEM from 3 experiments. **(D)** Cell cycle profile of individual p53-WT or KO RPE1-FUCCI cells to analyse multiple rounds of division following washout from 1, 4 or 7 days of treatment with palbociclib (1.25μM). Graph shows 50 cells analysed at random from one experiment, which is representative of 2 experimental repeats. **(E)** Quantification of cell cycle profiles from cells treated as in D. Graph shows 100 cells analysed at random from 2 experimental repeats.

To determine whether these cell cycle defects were associated with a reduction in long-term proliferation, we performed colony-forming assays under identical conditions. Figure 2C shows that 4-days of palbociclib treatment is sufficient to dramatically reduce colony forming potential in p53-WT cells, whereas 7 days of palbociclib is required to cause a similar reduction in p53-KO cells. We were struck by two major differences between the long-term proliferation data and the cell cycle analysis (Figures 2B,C). Firstly, 4-days of palbociclib treatment induced relatively few cell cycle withdrawals in p53-WT cells (16 % S/G2>G1 conversions) but this was associated with a strong reduction in long-term proliferation. Secondly, although removal of p53 allowed all cells to progress into mitosis following 4 or 7-day palbociclib treatment (Figure 2B, lower panels), p53 loss could only restore long-term proliferation in the 4-day treatment group (Figure 2C). Our FUCCI analysis only allowed quantification of the first cell cycle following drug release, because cells were released from palbociclib in the presence of the Eg5 inhibitor S-trityl-L-cysteine (STLC) to block cells in mitosis ^48^. To analyse additional cell cycles after release, we performed FUCCI analysis without STLC and analysed the first 3 days of proliferation following palbociclib release. This demonstrated that although most p53-WT and KO cells were able to complete the first cell cycle following washout from 4-day palbociclib treatment, only the p53-KO cells were able to continue to proliferate at a normal rate during subsequent cell cycles (Figure 2D and E), consistent with the difference in colony forming potential observed in the 4-day treatment groups (Figure 2C). The proliferative ability of p53-KO cells was compromised after 7 days of palbociclib treatment however, since considerably fewer mitotic events were apparent during the first 3 days following drug washout. This pattern also correlated with the reduction in long-term growth in this condition (Figure 2C). In general, cell cycle behaviour over the first 3 days was predictive of long-term growth potential, with only the normally dividing cells (i.e. approx. 24h cell cycles) able to form visible colonies at 10 days (Figure 2C,D). Therefore, CDK4/6 inhibition for longer than 3 days causes defects in subsequent cell cycles which restricts long-term proliferative potential. This effect can be partially rescued by knockout of p53 which allows cells to tolerate an extended window of palbociclib treatment before they begin to exit the cell cycle. This may explain why p53 loss is associated with resistance to CDK4/6 inhibition in patients ^16,49^.

We next investigated the reason for cell cycle withdrawal following CDK4/6 inhibition. The ability of p53 to induce cell cycle exit from G2 has previously been linked to p21 induction ^41,47^, therefore we analysed p21 levels following CDK4/6 inhibition. In p53-WT cells, we observed a strong induction of p21 after the release from prolonged CDK4/6 inhibition (Figure 3A,B). This p21 induction was absent in p53-KO cells (Figure 3B), as expected, which is consistent with the inability of these cells to exit the cell cycle from G2 (Figure 2B). The lack of p53-induced p21 had dramatic consequences, because instead of withdrawing from the cell cycle, p53-KO cells underwent a catastrophic mitosis that produced excessive DNA damage, as judged by nuclear morphology and γH2AX staining (Figure 3C-E). This also caused the appearance of symmetrical 53BP1 nuclear bodies after mitosis, a phenotype that results from the segregation of incompletely replicated chromosomes (Figure 3F) ^50,51^. Live-cell imaging of GFP-53BP1/H2B-RFP p53-KO RPE1 cells confirmed that DNA damage specifically appeared after an abnormal mitosis, and this was frequently associated with the segregation of unaligned or lagging chromosomes (Figures 3G-I). Examples of the abnormal divisions can be seen in movie-S1 (7 days palbociclib washout) in comparison to movie-S2 (1 day palbociclib washout). Cells have intrinsic mechanisms to either replicate or resolve incompletely replicated DNA during mitosis, but these systems can be overwhelmed under conditions of replication stress to cause DNA strand breaks during mitosis ^52-56^. To examine whether DNA replication was indeed ongoing during mitosis we performed mitotic DNA replication assays by examining EdU incorporation in mitotic cells 15 min after release from RO-3306 washout. Aphidicolin treatment, a well-known inducer of replication stress, was sufficient to elevate the levels of mitotic DNA replication in p53-KO cells, as expected ^54^. This increase in mitotic DNA replication was also observed after release from a prolonged palbociclib arrest (Figure 3G,H), consistent with the notion that DNA replication is also perturbed in these cells. Note, very few p53-WT cells enter mitosis after prolonged palbociclib release, which we hypothesise is due to a combination of p53-dependent cell cycle withdrawal (Figure 2B) and an intact ATR-dependent checkpoint that prevents mitotic entry until DNA replication is complete ^57^.

**Figure 3.**
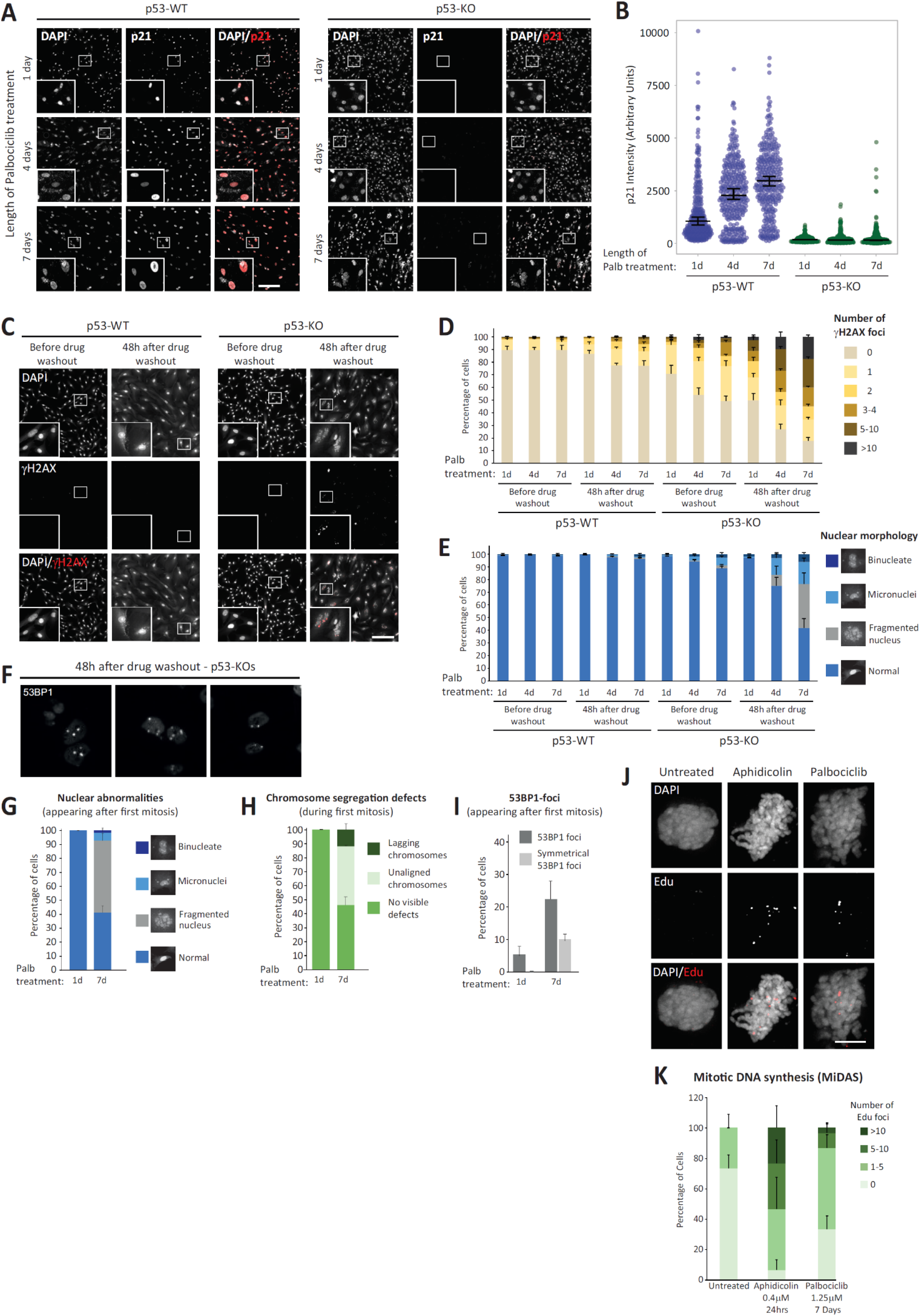
Prolonged CDK4/6 inhibition induces replication stress and p53-dependent cell cycle withdrawal. **(A)** Representative immunofluorescence images of p21 levels in p53-WT or KO RPE1 cells, 48 hours after release from 1, 4 or 7 days palbociclib (1.25μM) treatment. Zoom inserts are 3x magnification of the indicated regions. Scale bars = 250 μM. **(B)** Quantification of p21 intensities in cells treated as in panel A. At least 100 cells were analysed per experiment and graph shows data from 3 experimental repeats. Violin plots display the variation in intensities between individual cells. Horizontal lines display the median, and error bars show 95% confidence intervals. **(C)** Immunofluorescence images of DAPI and yH2AX staining in p53-WT or KO RPE1 cells either before or 48 hours after release from a 7-day treatment with palbociclib (1.25μM). Scale bar = 250 μM, zoom inserts = 3x magnification of highlighted regions. **(D)** Quantification of the γH2AX-positive DNA damage foci following palbociclib (1.25μM) treatment in p53 WT and KO RPE1 cells. Cells were treated for 1, 4 or 7 days and then analysed before or after drug washout for 48 hours. yH2AX foci were counted in 50 cells per condition per experiment and bar graphs represent mean data -/+ SEM from 6 experiments. **(E)** Quantification of the nuclear morphologies from cells treated as in panel D. 100 cells were scored per condition per experiment and bar graphs represent mean data -/+ SEM from 6 experiments. **(F)** Immunofluorescence images to demonstrate symmetrical 53BP1 staining following mitotic exit in p53-KO cells after release from 7 days of palbociclib arrest. 3 separate examples displayed. **(G)** Percentage of GFP-53BP1/H2B-RFP P53-KO RPE1 cells that display nuclear abnormalities specifically following the first mitosis after release from 1-day or 7-days palbociclib (1.25μM) treatment. >50 cells analysed from 2 experiments. **(H)** Percentage of GFP-53BP1/H2B-RFP P53-KO RPE1 cells with visible chromosome segregation defects during the first mitosis after release from 1-day or 7-days palbociclib (1.25μM) treatment. **(I)** Percentage of GFP-53BP1/H2B-RFP P53-KO RPE1 cells with visible 53BP1-foci appearing following the first mitosis after release from 1-day or 7-days palbociclib (1.25μM) treatment. **(J)** Representative immunofluorescence images of mitotic DNA replication assays (MiDAS). EdU foci in Nocodazole arrested p53 KO RPE1 cells following release from 7 days of palbociclib (1.25μM). Scale bar= 5 μM, zoom inserts = 3x magnification of highlighted areas. **(K)** Quantification of EdU foci in p53 KO RPE1 cells treated as in panel J. 10 cells were analysed per experiment and the stacked bar chart shows the mean -/+ SEM from 3 experimental repeats.

In summary, a prolonged palbociclib arrest causes replication stress following release from that arrest. This inhibits long-term viability by either inducing a p53-dependent withdrawal from the cell cycle or, in the absence of p53, by causing cells to undergo a catastrophic mitosis resulting in DNA damage as under-replicated chromosomes are segregated. If this damage is too excessive then long-term proliferation is still affected, as observed following release from a 7-day palbociclib arrest (Figures 2C, 2D and 3C-E). If the replication stress is milder, for example following 4-day palbociclib arrest, then cells can progress through mitosis but frequently arrest in a p53-dependent manner in the subsequent G1 (Figure 2B-D). This is consistent with the previous observations that mild replication stress cause a p21-dependent arrest in the subsequent G1 ^58,59^.

Defects in the cell cycle begin to appear if CDK4/6 inhibitors are applied for longer than 2 days (Figure 1 and S2). Therefore, to screen for potential causes of replication stress we performed a proteomic comparison of cells arrested in palbociclib for 2 or 7 days (Supplementary dataset 1). Out of the top 15 most significantly changing proteins, 5 were members of the MCM2-7 complex, which licences DNA replication origins and then forms the catalytic core of the CMG (Cdc45-MCM2-7-GINS) helicase that is responsible for unwinding DNA to allow replication fork progression (Figure 4A,B) ^60^. In addition to MCMs, many other components of the core replisome were downregulated by prolonged palbociclib treatment, including the DNA clamp (PCNA), the clamp-loading complex (RFC1-5) and many accessory factors that bind PCNA (FEN1, DNMT1, FAM111A). In addition, we observed downregulation of a variety of DNA polymerases along with their accessory subunits (Figure 4C). Western blotting confirmed that the levels of replisome components progressively decreased during a palbociclib arrest and, importantly, remained low after palbociclib washout for 8 or 24h (Figure 4D,E); timepoints chosen to capture the majority of cells as they replicate DNA in S-phase (Figure 1C). In addition to decreasing total MCM protein, palbociclib treatment also reduced the extent of origin licencing after release from the inhibitor, as assessed by the level of chromatin-bound MCM during early S-phase (Figure 4F). Therefore, a palbociclib-induced G1 arrest decreases the level of replisome components and reduces the number of licenced replication origins available during S-phase. This likely explains why if the G1-arrest is too long, there is a failure in DNA replication after release from that arrest, resulting in either cell cycle exit (p53 proficient) or a catastrophic mitosis with under-replicated DNA (p53 deficient). Note that the decrease in replisome components and origin licencing was similar in p53-KO cells (Figure 4D-F and S5), implying that p53 status primarily defines the response to DNA replication defects.

**Figure 4.**
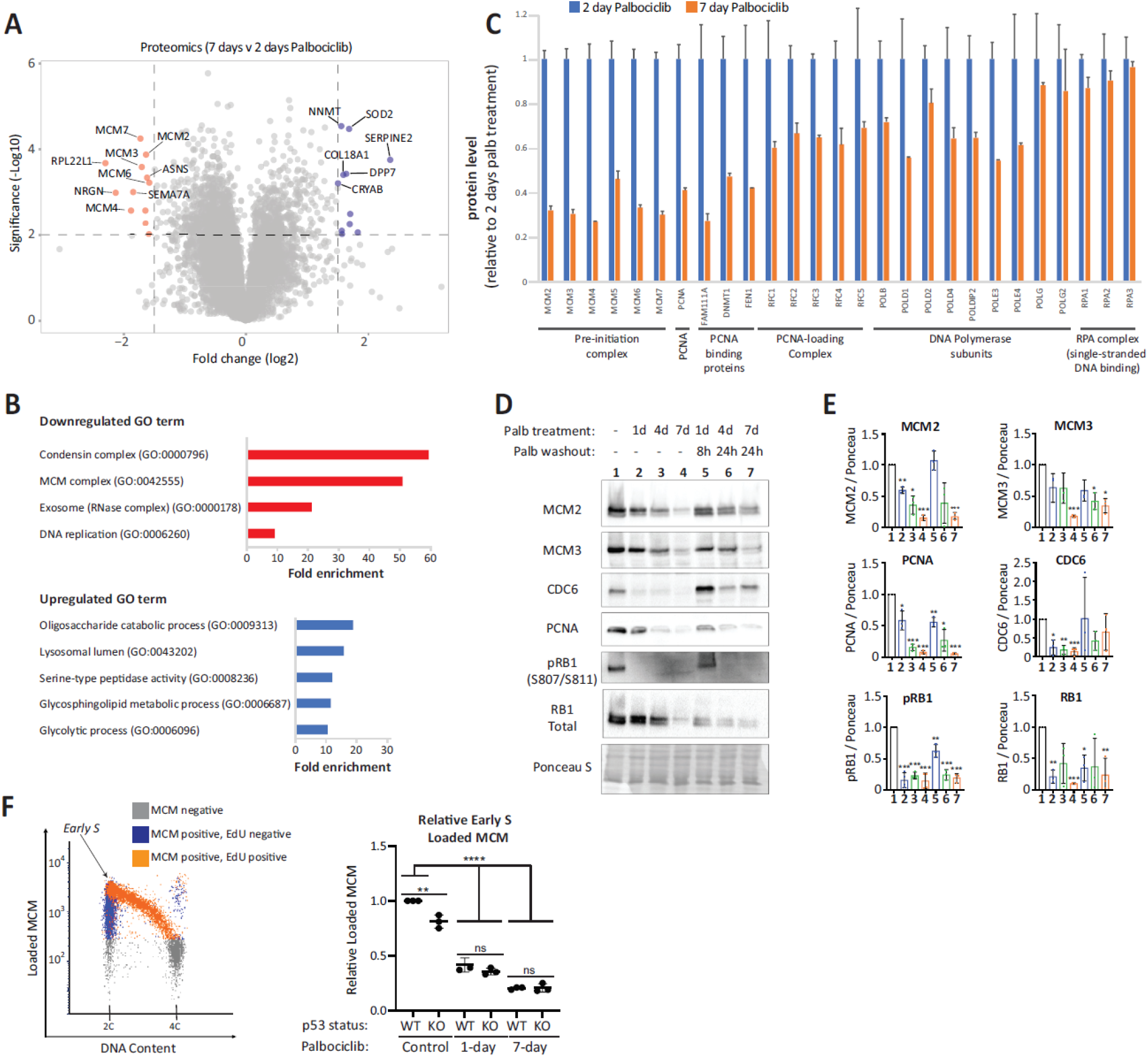
Prolonged G1 arrest following palbociclib treatment downregulates replisome components and impairs origin licencing. **(A)** Volcano plot of proteins up or downregulated following prolonged palbociclib (1.25μM) treatment in RPE1 cells. The top 10 significantly upregulated and downregulated proteins are shown in blue and red, respectively. **(B)** The top up or downregulated Gene Ontology (GO) terms following 7-day palbociclib (1.25μM) treatment relative to 2 days of treatment. **(C)** Quantification of relative change in protein levels of selected replisome components between 2-day (blue bars) and 7-day (orange bars) palbociclib (1.25μM) treatment. **(D)** Representative western blots of whole cell lysates from RPE1-WT cells treated with palbociclib (1.25μM) for 1, 4 or 7 days, or treated identically, and then washed out for the indicated times to reflect when the majority of cells are in S-phase (see Figure 1C). **(E)** Analysis of adjusted relative density from 3 independent western blot experiments. Bars display mean values -/+ SD. Significance determined by unpaired Student’s T test comparing treated target protein to asynchronous target control. (* < 0.01, ** < 0.001, *** < 0.0001). **(F)** The left panel shows a representative plot of MCM loaded in untreated RPE1 cells used to generate the corresponding graph at right. Soluble MCM was pre-extracted from cells and the amount of the remaining DNA-loaded MCM was analysed by flow cytometry. DNA content was measured with DAPI, and DNA synthesis was measured using a 30-minute EdU pulse. The population of early S phase cells (2C DNA content, EdU positive) analysed indicated. RPE1 cells were treated with palbociclib (1.25uM) for 1 or 7 days followed by drug washout for 8 hrs after 1 day of arrest or 24 hrs after 7 days of arrest to capture early S phase. The amount of DNA-loaded MCM in early S phase cells was compared to untreated control cells. The measured fluorescent intensity of each sample was divided by the background intensity of an identically treated but unstained control. The resulting ratios were normalized to WT control cells. p53 status is as indicated. Significance determined by one-way ANOVA followed by Tukey’s multiple comparisons test (** p = 0.001, **** p < 0.0001).

The ability of CDK4/6 inhibitors to induce genotoxic damage as a result of replication stress has important implications for cancer treatment. Firstly, it suggests that tumour cells with ongoing replication stress maybe more sensitive to the long-term effects of CDK4/6 inhibition. Secondly, it implies that chemotherapeutics that enhance replication stress may sensitize cells to CDK4/6 inhibition. We sought to address these points in a controlled manner using RPE1 cells.

Aneuploidy, a well-established hallmark of tumourigenesis, is known to induce replication stress ^61-64^. We therefore induced widespread aneuploidy in the otherwise near-diploid RPE1 cells by treatment with 0.5μM reversine for 24h (an MPS1 inhibitor that induces aneuploidy by inhibiting the mitotic checkpoint: see ^61^). The majority of aneuploid RPE cells (>90%) were able to transit though S/G2 phase and enter mitosis similarly to their euploid parental cells, as shown previously ^61^ (Figure 5A,B). However, following a palbociclib arrest, aneuploid cells exhibited many more cell cycle withdrawals (Figure 5A vs 5B) and displayed a reduced ability to form colonies (Figurer 5G). This demonstrates that aneuploidy is sufficient to induce vulnerability to CDK4/6 inhibition, perhaps by elevating the levels of endogenous replication stress ^61-64^.

**Figure 5.**
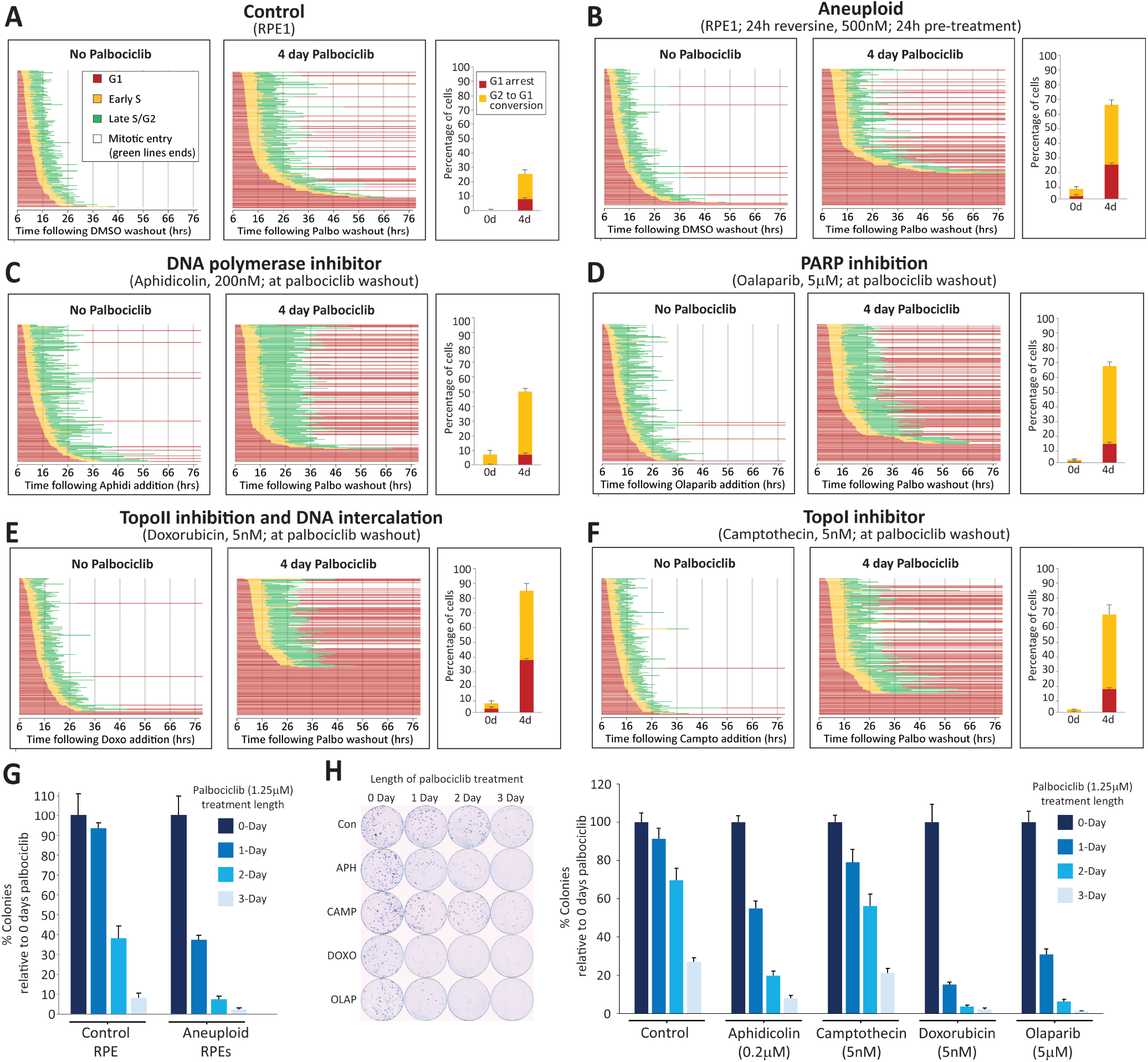
Genotoxic chemotherapeutics sensitize RPE1 cells to CDK4/6 inhibition. **(A)** Cell cycle profile of individual RPE1-FUCCI cells (each bar represents one cell) after release from 4d treatment with palbociclib (1.25μM) or DMSO. **(B)** Cell cycle profile of aneuploid RPE1-FUCCI, treated as in panel A. To generate aneuploidy, RPE1 cells were treated with 0.5μM reversine for 24h immediately prior to palbociclib treatment, as described previously ^61^. **(C-F)** Cell cycle profile of individual RPE1-FUCCI cells treated as in panel A, but additionally treated after drug washout with aphidicolin (C), olaparib (D), doxorubicin (E) or camptothecin (F), at indicated concentrations. **(G)** Colony forming assays with control or aneuploid RPE1 cells treated with palbociclib (1.25μM) for indicated times and then grown at low density without inhibitor for 10 days. Aneuploid cells were created as stated in panel B. Each bar displays mean data -/+ SD from 3 experiments. **(H)** Colony forming assays with RPE1 cells treated with palbociclib (1.25μM) for indicated times, and then grown at low density without palbociclib for 10 days. DMSO (control) or indicated genotoxic drugs were applied for the first 24 h after palbociclib washout. Each bar displays mean data -/+ SD from 4 experiments. All experiments in panels A-F where run at the same time to allow comparison to the same control (panel A). STLC (10 μM) was added in all movies to prevent progression past the first mitosis. In all FUCCI graphs, 50 cells were analysed at random for each repeat and 3 experimental repeats are displayed (150 cells total).

We next used a low dose of aphidicolin to partially inhibit DNA polymerases and induce replication stress directly. Figure 5C shows that whilst this did not have a strong effect on cell cycle progression when given alone, it was able to enhance the number of cell cycle withdrawals when given to cells released from CDK4/6 inhibition. A number of genotoxic anti-cancer drugs also induce replication stress, therefore we analysed the effect of three such compounds that impede DNA replication differently: camptothecin (TopoI inhibitor), doxorubicin (TopoII inhibitor) or olaparib (PARP inhibitor). We chose a dose of each drug previously shown to be sublethal in RPE1 cells ^65^, and demonstrated that this produced only mild effects on cell cycle timing and progression in control cells (Figure 5D-F). However, when given following a palbociclib arrest, these drugs caused the majority of cells to fail to complete the first cell cycle. In particular, there was a large increase in cells that commenced DNA replication but then withdrew into G1 before entering mitosis. Colony forming assays demonstrated a strong sensitization between genotoxic chemotherapeutics and CDK4/6 inhibition, with doxorubicin and olaparib causing a strong reduction in proliferation after only 1-day of palbociclib treatment (Figure 5H). These data suggest that combining CDK4/6 inhibition with existing genotoxic drugs may be a promising therapeutic strategy, as also suggested recently by others, but for different reasons ^66^ (see discussion).

## Discussion

A major finding of this study is that CDK4/6 inhibitors, like many other broad-spectrum anti-cancer drugs, induce genotoxic stress during S-phase. This may initially appear counterintuitive for a class of drugs that effectively prevent S-phase entry. However, when cells are arrested in G1 following CDK4/6 inhibition, key components of the replisome are downregulated and, if this is allowed to proceed for too long, then DNA replication is perturbed upon release from that G1 arrest. Therefore, we propose that long-term toxicity following CDK4/6 inhibition depends on at least two key factors: 1) the duration of time that cells remain arrested in G1, and 2) how well cells can then tolerate the resulting replication stress.

Using a non-transformed near-diploid RPE1 cell line, we demonstrate that the G1-arrest becomes problematic if it lasts for longer than 2 days. This timing is dependent on the level of endogenous replication stress, because if this is elevated pharmacologically then RPE1 cells withdraw from the cell cycle more readily and they are sensitive to shorter durations of palbociclib treatment (Figure 5A-H). Therefore, we hypothesise that tumour cells, many of which have elevated levels of replication stress ^67^, will also be sensitive to a short G1 arrest. We addressed this concept in a controlled manner in RPE1 cells by inducing aneuploidy; a hallmark of most cancer cells that is known to cause replication stress ^61-64^. This demonstrates that aneuploid RPE1 cells exhibit more cell cycle withdrawals following palbociclib treatment (Figure 5A,B) and the length of G1 arrest that they can tolerate is approximately 1-day shorter than their euploid counterparts (Figure 5G). Therefore, a common feature of tumourigenesis, which increases replication stress, also induces sensitivity to a prolonged G1-arrest. Tumour cells frequently contain mutations that promote an efficient G1-arrest following CDK4/6 inhibition ^21^, therefore this “double hit” - a longer G1 arrest and a lower tolerance to that arrest - could explain their exquisite sensitivity.

It is important to state that the RPE1 cells we use in this study, although telomerase-immortalised and non-transformed ^68^, do still have mutations in at least two known cancer-associated genes: *CDKN2A* and *KRAS* ^69,70^. *CDKN2A* deletion causes sensitivity to palbociclib ^71-74^, therefore the CDKN2A mutation may explain why RPE1 cells arrest so efficiently in G1 following CDK4/6 inhibition (the IC50s for palbociclib and abemaciclib are comparable to the most sensitive cancer cell type in a panel of 560 tested lines: see Figure 1b and ^21^). It is also possible that the activating *KRAS* mutation ^69^ contributes to the phenotype of CDK4/6 inhibition; for example, by causing oncogene-induced replication stress ^75^. It is therefore important to build on this work in future to examine the response of a large panel of cell types (non-transformed and tumour lines) to different lengths of G1-arrest. This information will help to identify genetic backgrounds that sensitise cells to the effects of this arrest and may reveal biomarkers that can predict long-term response. We hypothesise that candidates in this regard will fall into at least three categories: 1) factors that determine the initial response in G1, for example, by controlling the downregulation of replisome components; 2) factors that control how well the resulting stress during S-phase is tolerated, for example, mutants that induce replication stress or inhibit the ability of cells to repair replication defects; and 3) downstream mediators that determine the fate of cells following this genotoxic stress. Our data demonstrates that p53 is a critical downstream fate determinant that controls cell cycle withdrawal, and this may ultimately help to explain why p53 loss is associated with resistance to CDK4/6 inhibition in the clinic ^16,49^. To facilitate future efforts to determine sensitive/resistant cell types, it will also be critical to understand why replisome components are downregulated during a prolonged G1-arrest (Figures 3A-D) and to determine why the extent of origin licencing is significantly impaired following just 1-day of palbociclib treatment (Figure 3F).

An important finding of this work is that CDK4/6 inhibition sensitizes cells to cytotoxic chemotherapeutics currently in widespread clinical use. This was also demonstrated recently in pancreatic ductal adenocarcinoma (PDAC); a tumour type that is similarly characterised by activating KRAS(G12V) and CDKN2A mutations ^66^. In this case, the sensitivity in PDAC models was attributed to the ability of palbociclib to prevent DNA repair. It is possible that Cdk4/6 inhibition promotes genotoxic damage by elevating replication stress and also inhibiting DNA repair, however, it is important to carefully distinguish between these possibilities because it may determine the optimal order of drug exposure. It is worth noting in this regard, that the application of anti-mitotic drugs (monastrol, BI2536 or paclitaxel) sensitized various PDAC models to a subsequent palbociclib arrest ^66^. These drugs all induce aneuploidy; therefore, we speculate that at least part of this sensitization is due to elevated levels of replication stress. When assessing optimal cytotoxic combinations in preclinical models, it will be important to assess if drug-free periods following CDK4/6 inhibition elevate DNA damage by allowing progression through S-phase.

In clinical practice, palbociclib and ribociclib are typically given in cycles of 3-weeks on, 1-week off; primarily to allow recovery from haematopoietic toxicity ^76^. It is possible that these drug-holiday periods contribute to tumour cell killing by allowing replication stress to cause DNA damage. It is important to test this hypothesis because, if elevated DNA damage is detected when CDK4/6 inhibitors are withdrawn, then the timing/duration of drug holidays could potentially be optimised. On this point, it should be noted that abemaciclib is dosed continuously due to a low level of haematopoietic toxicity ^76^. However, this does not exclude the possibility that replication stress occurs in this situation as well. That is because sub-maximal inhibition of CDK4/6 extends G1 duration in RPE1 cells (Figure S6) ^77^. Therefore, cells continually exposed to the right concentration of drug may experience recurrent delays in G1 prior to S-phase entry, which could be sufficient to induce cycles of replication stress, as long G1 length is extended sufficiently.

Finally, CDK4/6 inhibition can re-sensitize tumours to immune checkpoint blockade ^25-31^. The ability of CDK4/6 inhibition to inhibit DNA replication could help to promote immune engagement in a number of different ways. Replication stress can activate the cGAS/STING-mediated interferon (IFN) response and increase the number of mutations/neoantigens ^78,79^. In addition, it can induce senescence, thereby creating non-proliferative tumour cells that continually secrete factors to engage the immune system ^25-28^. CDK4/6 inhibitor-induced senescence helps to sensitize tumours to immune checkpoint blockade ^29-31^, therefore it will be important to test whether replication stress leads to senescence in these settings. There are currently 14 clinical trials ongoing in 8 tumour types to assess whether CDK4/6 inhibition can improve response to anti-cancer immunotherapy ^32^, therefore it will also be important to assess whether p53-status correlates with response in these situations. P53 may behave like a double-edged sword in this regard, because although it is typically required for entry into senescence ^33^, if p53 is absent, then severely under-replicated chromosomes are allowed to progress into mitosis (Figure 3) and the resulting micronuclei can activate the cGAS-STING pathway ^80-83^.

In summary, the work presented here links CDK4/6 inhibitors with genotoxic stress, which now provides a rationale to better understand how these drugs might selectively target tumour cells. CDK4/6 inhibitors are already known to arrest tumour cells more efficiently in G1 ^2,21^, but if they also capitalise on the fact that these tumours are exquisitely sensitive to that arrest as a result of ongoing replication stress, then the implications for cancer treatment could be wide-ranging. It is therefore now critical to build on this work and carefully examine these concepts in preclinical and clinical settings to determine whether replication stress is a common outcome of CDK4/6 inhibition.

## Supporting information

Movie-S1: 7-day palbo washout

Movie-S2: 1-day palbo washout

## Acknowledgements

We thank staff at the Dundee Imaging Facility and the Genetic Core Services Unit. This work was supported by the Medical Research Council (studentship to L.C.), Tenovus Scotland (studentship for R.F.), a Cancer Research UK Programme Foundation Award to A.T.S (C47320/A21229. which also funds A.C.), a Wellcome/Royal Society Sir Henry Dale Fellowship to T.L. (206211/Z/17/Z) and the United States National Institutes of Health grants (GM083024 and GM102413 to J.G.C.; B.L.M was supported by T32GM135128, and R.H.W. was supported by T32CA009156). The WCB Proteomics core is supported by Wellcome (108504 and 203149). The UNC Flow Cytometry Core Facility is supported in part by a National Institutes of Health Cancer Core Support Grant to the UNC Lineberger Comprehensive Cancer Center (CA016086).

## Contributions statement

A.T.S. conceived the study and supervised L.C. and R.F. who performed the majority of experiments. T.L. designed and supervised the MS analysis that first identified reduced MCM levels, with A.C. and C.S. providing help with LC-MS/MS sample preparation and analysis. J.G.C. designed and supervised experiments to characterise MCM loading defects, with B.L.M performing the MCM FACS analysis and R.H.W carrying out Western analysis to quantify replisome components. A.T.S. wrote the manuscript with comments from all authors.

## materials and methods

### Cell culture and reagents

hTERT-RPE1 (RPE1) were purchased from ATCC and the RPE1-FUCCI were published previously ^41^. Cells were cultured at 37°C with 5% CO_2_ in DMEM supplemented with 9% FBS and 50ug/ml penicillin/streptomycin. All cells were authenticated by STR profiling (Eurofins) and screened for mycoplasma every 1-2 months. The following drugs were used in this study: Palbociclib (PD-0332991, hydrochloride salt, MedChemExpress, HY-50767A); ribociclib (LEE-011, Selleckchem, S7440); abemaciclib (LY-2835219, Selleckchem, S7158); trilaciclib (G1T28, Insight, HY-101467A); S-Trityl-L-cysteine (STLC, Sigma Aldrich, 167739), aphidicolin (Santa Cruz, SC-201535), reversine (Sigma, R3904), doxorubicin (Selleckchem, S1208), olaparib (Selleckchem, S1060), camptothecin (Sigma, C9911), nutlin-3a (Sigma, SML0580), DAPI (4’,6-Diamidino-2-Phenylindole; Thermo Fischer, D1306), RO3306 (Tocris, #4181), EdU (Sigma-Aldrich, BCK-EDU488), nocodazole (Sigma-Aldrich, #487928).

### Immunofluorescence

Cells were plated on High Precision 1.5H 12-mm coverslips (marienfeld) and fixed for 10 minutes with 4% paraformaldehyde dissolved in PBS. Once fixed, coverslips were washed three times in PBS and then blocked in 3% BSA dissolved in PBS with 0.5% Triton X-100 for 30 minutes. Coverslips were then incubated with primary antibodies at 4°C overnight, prior to washing with PBS and incubation with secondary antibodies and DAPI (1μg/ml) for 2-4 hours at room temperature. After further washing, coverslips were mounted onto slides with ProLong Gold Antifade (Thermo Fischer, P10144). Coverslips were images on either a Zeiss Axio Observer using a Plan-apopchromat 20x/0.8 M27 Air objective or a Deltavision with a 100x/1.40 NA U Plan S Apochromat objective. The primary antibodies used were: rabbit anti-p21 (H-164) (Santa Cruz, sc-756; 1/500), mouse anti-p53 (Santa Cruz, sc-126; 1/1000), mouse anti-phospho-Histone H2A.X (Ser139) (Sigma, 05-636; 1/1000), rabbit anti-53BP1 (Novus biologicals, NB100-304; 1/1000). The secondary antibodies used were highly-cross absorbed goat anti-rabbit or anti-mouse coupled to Alexa Fluor 488, Alexa Fluor 568 which were all used at 1/1000 dilution. All antibodies were made up in 3% BSA in PBS. For Edu staining, a base click EdU staining kit was used (Sigma, BCK-EDU488), as per manufacturer’s instructions.

### MiDAS Protocol

RPE1 p53 KO cells were plated at low confluence in 10cm dishes and treated with palbociclib (1.25uM) for 7 days. Palbociclib was then removed via an extensive washout and cells were transferred coverslips. Coverslips were then returned to incubation for 16hrs before being treated with RO3306 (10µMs) for a further 2 hours to enrich for cells in G2. Media was then exchanged twice over 15minutes and cells were treated with EdU (10μM) and nocodazole (3.3μM). After 1-hour cells were fixed in 4% PFA. Note that control cells were either treated with aphidicolin (0.4μM) for 40 hours or left untreated prior to addition of RO3306. Following fixation cells were permeabilised with 0.2% Triton-X in 3% BSA dissolved in PBS. Staining of incorporated EdU was carried out as per manufacturers’ instructions and coverslips were mounted onto slides using prolong gold antifade. Coverslips were imaged using a Deltavision with a 100x/1.40 NA U Plan S Apochromat objective.

### Time-lapse imaging

For FUCCI time-lapse imaging, cells were imaged in 24-well plates in DMEM inside a heated 37°C chamber with 5% CO2. Images were taken every 10 minutes with a 10x/0.5 NA air objective using a Zeiss Axio Observer 7 with a CMOS ORCA flash 4.0 camera at 4×4 binning. GFP-53BP1/H2B-RFP RPE1 cell lines were imaged on a DeltaVision Elite system in 8 well chambers (Ibidi) in L15 media within a heated 37°C chamber. Images were taken every 4 minutes with a 40x/1.3 NA oil objective using a DV Elite system equipped with Photometrics CascadeII:1024 EMCCD camera at 4×4 binning.

### Generating knockout cell lines

To generate p53 knockout cells, a gRNA targeting exon 4 of p53 (ACCAGCAGCTCCTACACCGG) was cloned into the pEs-gRNA vector by site directed mutagenesis, as described previously ^84^. RPE1 and RPE1-FUCCI cells were then transfected with this gRNA vector along with a pcDNA5-Cas9 vector in a 3:1 ratio. Knockout cells were subsequently selected by cultured in 5μM of Nutlin-3a until no visible cells remained on the control non-transfected plates (approximately 3 weeks). P53 knockout status was confirmed via immunoblotting and immunofluorescence.

### Western blotting

Total protein lysates for immunoblot were prepared by harvesting cells in trypsin, pelleting, and flash freezing. Cell pellets were lysed in ice cold RIPA buffer (50 mM Tris ph 7.6, 150 mM NaCL, 1% NP40, 0.5% sodium deoxycholate, 1mM EDTA, 0.1% SDS and protease and phosphatase inhibitors (0.1 mM Pefabloc, 1 μg/ml pepstatin A, 1 μg/ml leupeptin, 1 μg/ml aprotinin, 10 μg/ml phosvitin, 1 mM β-glycerol phosphate, and 1 mM sodium orthovanadate) on ice for 20 min. Lysates were centrifuged at 13,000 g and 4°C for 10 min, followed by Bradford assay (Biorad) to determine equal amounts of protein to load per lane. Samples were mixed with loading buffer to final concentrations of: 1% SDS, 2.5% 2-mercaptoethanol, 0.1% bromophenol blue, 50 mM Tris, pH 6.8, and 10% glycerol. Samples were boiled, then separated on SDS-PAGE gels, and transferred polyvinylidene difluoride membranes (Thermo Fisher Scientific). After transfer, blots were blocked in 5% milk in TBS with 0.1% Tween 20 (TBS-T) and incubated overnight at 4°C in primary antibody in TBS-T. Then membranes were washed in TBS-T 3x, incubated in HRP-conjugated secondary antibody for 1 h at RT, washed in TBST 3x, and imaged with ECL Prime (Amersham). Membranes were stained with Ponceau S (Sigma-Aldrich) to visualize protein loading. Antibodies used were mouse anti-MCM2 (BM28) (BD Biosciences, 610701; 1/1000), rabbit anti-MCM3 (A300-192A, Bethyl Laboratories; 1/1000), mouse anti-CDC6 (180.2) (sc-9964, Santa Cruz; 1/500), mouse anti-PCNA (sc-25280, Santa Cruz; 1/1000), mouse anti-RB (554136, BD Biosciences), rabbit anti-pRB (Ser807/Ser811) (9308S, Cell Signaling Technology; 1/1000), rabbit anti-p21 (H-164) (Santa Cruz, sc-756; 1/250), mouse anti-p53 (DO-1) (Santa Cruz, sc-126; 1/1000), rabbit anti-actin (Sigma, A2066; 1/5000). Anti-Mouse HRP (1858413, Pierce; 1/10,000), and anti-Rabbit HRP (1858413, Pierce; 1/10,000) secondary were used.

### Chromatin-bound MCM FACS assays

The amount of DNA-loaded MCM following release from palbociclib treatment was analysed as described previously ^85^. RPE1 WT or p53 KO cells were treated with palbociclib for 1 or 7 days and the drug was washed out for 8 or 24 hours, respectively. 30 minutes prior to cell collection, cells were pulse labelled with 10 μM EdU (Sigma) to monitor DNA synthesis. Soluble MCM was pre-extracted from cells on ice for 10 minutes in cold CSK buffer (10 mM Pipes pH 7.0, 300 mM sucrose, 100 mM NaCl, 3 mM MgCl_2_) supplemented with 0.5% triton X-100, protease inhibitors (0.1 mM AEBSF, 1 μg/mL pepstatin A, 1 μg/mL aprotinin, 0.1 mM PMSF), and phosphatase inhibitors (10 μg/mL phosvitin, 1 mM β-glycerol phosphate, 1 mM Na-orthovanadate). After washing cells in PBS + 1% BSA, cells were fixed in PBS + 4% paraformaldehyde (Sigma) for 15 minutes at room temperature. Cells were processed for EdU conjugation to Alexa Fluor 647-azide (Life Technologies) by incubation in PBS containing 1 mM CuSO_4_, 1 mM AF-647, and 100 mM fresh ascorbic acid for 30 minutes at room temperature in the dark. The levels of DNA-loaded MCM were detected by incubating cells in anti-MCM2 BM28 antibody (1:200, BD biosciences, 610700) for 1 hr at 37°C in the dark followed by incubation in anti-mouse-488 secondary antibody (1:1000) for 1 hr at 37°C in the dark. DNA content was measured by incubating cells in 1 μg/mL DAPI and 100 μg/mL RNAase for 1 hr at 37°C in the dark or alternatively overnight at 4°C. Cells were analysed using an Attune NxT flow cytometer (Thermo Fisher Scientific) and data were analysed using FCS Express 7 Research (De Novo Software).

For each experimental condition, an identically treated sample was included but was not labelled for EdU or MCM in order to determine the limit of detection. Early S phase cells were analysed by gating on cells that had 2C DNA content and were EdU positive. For each sample, the mean AF-647 fluorescent intensity of early S phase cells was divided by the mean AF-647 fluorescent intensity of the identically treated but unstained control sample. The displayed data are the normalization of these ratios to asynchronous control cells.

### Colony forming assay

For the colony forming assays, cells were treated with palbociclib (1.25µM), ribociclib (5µM), abemaciclib (600nM), or trilaciclib (600nM) at 200,000 cells per 15cm dish for different length of time (1 – 7 days) prior to drug washout (6 x 1hr washes). Following washing cells were trypsined and plated in triplicate at 250 cells into 10cm dishes and left to grow for 10 days. For the experiments in figure 5, different genotoxic drugs were added for the first 24 hours after replating, before washout and incubation in standard media for the remaining 9 days. At the end of the assay, cells were washed twice in PBS and then fixed at 100% ethanol for 5 mins. Developing solution (1:1 ratio of 2% Borax:2% Toluene-D in water) was added to the fixed cells for 5 minutes and the plates were then rinsed thoroughly with water and left to dry overnight. The plates were then scanned and the number of colonies quantified using ImageJ. This was performed by cropping to an individual plate and converting to a binary image. The fill holes, watershed, and analyze particles functions were then used to count colonies.

### Mass spec sample preparation

Cells were plated in 15cm dishes and treated with palbociclib for 2 or 7 days. Cells were lysed in cell extraction buffer containing 2% SDS, 1X PhosStop (Roche) and 1x cOmplete protease inhibitor cocktail (Roche). An aliquot of extract containing 100 µg protein was then digested by benzonase (Merck) and precipitated by acetone. The protein pellet was resuspended in digest buffer (0.1 M triethylammonium bicarbonate, pH 8.5, Sigma-Aldrich, tandem mass tag (TMT) labeling using a 6-plex TMT kit (Thermo Fisher Scientific) and desalted. Peptides were then separated using high pH reverse phase chromatography (Waters BEH 4.6 mm×150 mm C18 column; A, 10 mM ammonium formate, pH 9.0; B, 80% acetonitrile plus 10 mM ammonium formate, pH 9.0) into 16 fractions ^86^. Fractions were then dried under vacuum and resuspended in 5% formic acid for liquid chromatography tandem mass spectrometry (LC-MS/MS) analysis.

### LC-MS/MS

LC-MS analysis was performed on an Orbitrap Fusion Lumos Tribrid MS (Thermo Fisher Scientific) coupled on-line, to an Ultimate 3000 RSLCnano HPLC (Dionex, Thermo Fisher Scientific). Peptides were separated on a 50 cm EASY-Spray column (Thermo Fisher Scientific) and ionized using an EASY-Spray source (Thermo Fisher Scientific) operated at a constant temperature of 50°C. Mobile phase A consisted of 0.1% formic acid in water while mobile phase B consisted of 80% acetonitrile and 0.1% formic acid. Peptides were loaded onto the column at a flow rate of 0.3 μl/min and eluted at a flow rate of 0.25 μl/min according to the following gradient: 2 to 40% mobile phase B in 120 min, then to 95% in 11 min. The percentage of mobile phase B remained constant for 10 min and returned to 2% until the end of the run (160 min).

MS1 survey scans were performed at 120,000 resolution (scan range 350–1500 *m*/*z*) with an ion target of 2.0×10^5^ and maximum injection time of 50 ms. For MS2, precursors selected using a quadrupole isolation window of 1.2 Th with an AGC target of 1E5 and a maximum injection time of 100 ms. Product ions from HCD fragmentation (32% normalised collision energy were then scanned using the Orbitrap with 30k resolution. Only ions with charge between 2 and 7 were selected for MS2.

### MS data analysis

Raw data files were processed using MaxQuant version 1.6.2.6 ^87^, which incorporates the Andromeda search engine ^88^. The spectra were searched against a human FASTA database (accessed June 2018) containing all reviewed entries in the reference UniProt Human Proteome. The processed output was then analyzed using R or RStudio software.

### Image quantification

To calculate the percentage of G1 arrested cells, RPE1-FUCCI cells were treated (as described in the legends) and then imaged using a Zeiss Axio Observer 7 with 10x/0.5 NA air objective and a CMOS ORCA flash 4.0 camera at 4×4 binning. Five positions were imaged per well using filtersets to image mKO2-cdt1 (red) and mAG-geminin (green). The TrackMate function in ImageJ was then used to quantify the number of RPE1-FUCCI cells in each channel. The percentage of red (G1-arrested) cells was calculated and used to generate dose-response curves in GraphPad Prism 7.

The single cells FUCCI profiles were generated manually by analysing RPE1-FUCCI movies. 50 red cells were selected and marked at random at the beginning of the movie. The time points in which the FUCCI cells change colour was recorded to determine the time spent in each phase of the first cell cycle following release from CDK4/6 inhibition. All images were placed on the same scale prior to analysis to ensure that the red/yellow/green cut-offs were reproducibly calculated between experiments. Mitotic entry was timed based on visualisation of typical mitotic cell rounding and loss of nuclear mAG-geminin signal.

P21 intensities were calculated using ImageJ and the first 100 cells in each image. The DAPI channel was used to generate an ROI overlay which was then applied to the p21 channel. The mean grey value of each ROI in the p21 channel was then measured along with the background intensity which was then subtracted from each of these values.

yH2AX foci were counted by eye in the first 50 cells (per condition) selected using the DAPI channel. For scoring of nuclear abnormalities, the first 100 cells within the image were counted and scored based on their nuclear morphology. 53bp1-H2B movies were analysed by eye quantifying nuclear morphologies as mentioned above. Chromosome alignment was also scored in cells that displayed H2B expression

## Supplementary legends

**Movie S1: First division after washout from 7-day palbociclib washout in p53-KO cells**. Frames taken every 4 mins to capture first division after palbociclib (7 days, 1.25μM) washout. Note, p53-KO cells are shown because p53-WT cells arrest prior to division and withdraw from the cell cycle.

**Movie S2: First division after washout from 1-day palbociclib washout in p53-KO cells**. Frames taken every 4 mins to capture first division after palbociclib (1 days, 1.25μM) washout.

**Supplementary Dataset 1: Proteomic Analysis of RPE1 cells arrest in palbociclib for 2 days or 7 days**. Cells treated with 1.25μM palbociclib for indicated times.

