## Supplementary Figures for "CDK4/6 inhibitors induce replication stress to cause long-term cell cycle withdrawal"

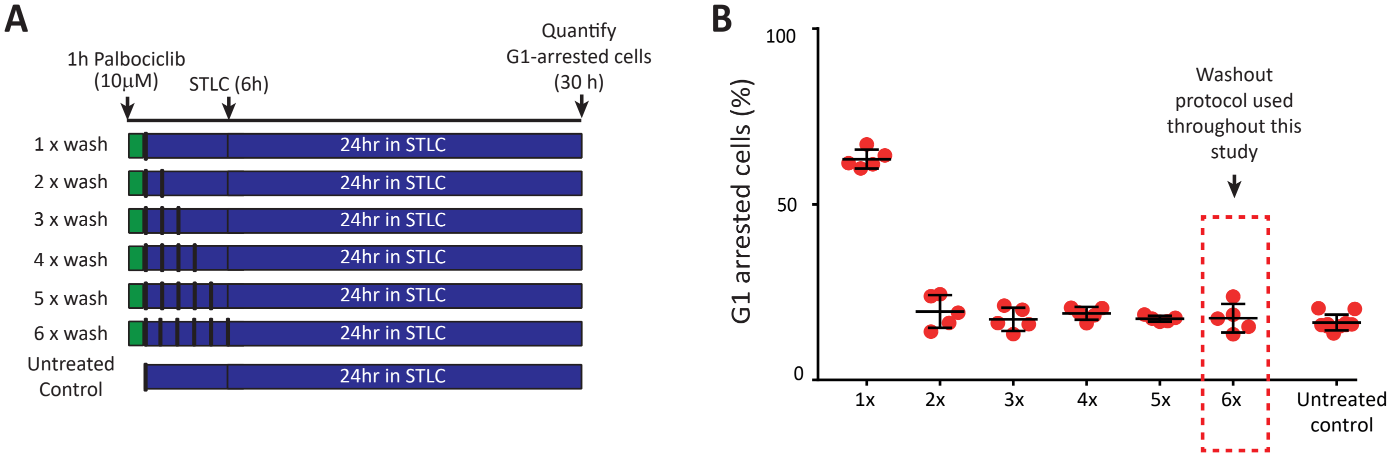


**Figure S1: Extensive drug-washout protocol used throughout this study. (A)** Schematic showing different washout protocols tested to ensure washout of high dose palbociclib. RPE1-FUCCI cells were treated for 1 hour with 10 µM palbociclib and subsequently washed out 1-6 times, with 1h equilibration periods interspersed between washes. STLC (10 µM) was then added to arrest cells in mitosis before quantifying the amount of red, G1-arrested cells 24 hours later. **(B)** Quantification of the G1-arrested cells following the washout protocol described in A. Graphs display mean data -/+ SD from 2 experiment, with at least 500 cells counted per condition per experiment. (The points represent 5 different positions that were imaged per condition). This data is representative of two experimental repeats.

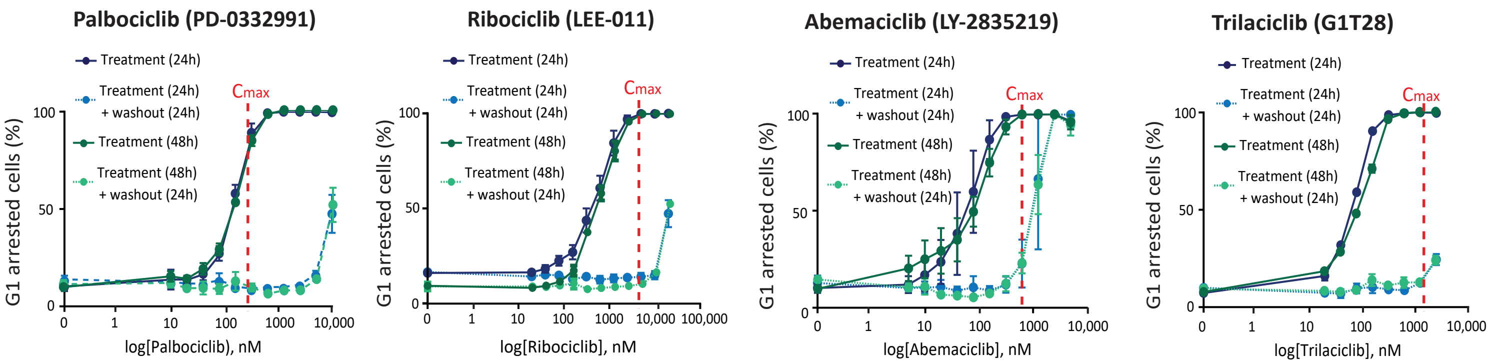


**Figure S2: 24 or 48 hours of CDK4/6 inhibition give similar dose-response and reversibility curves.** Percentage of G1-arrested RPE1-FUCCI cells after treatment for 48h with different CDK4/6 inhibitors (dark green solid lines) or 24h after subsequent drug washout (light green dotted lines). The data is overlaid with 24h arrest data from figure 1A (blue lines) to allow comparison. Vertical red dotted lines indicate Cmax values observed in patients (taken from 17, 40). Graphs display mean data -/+ SEM from 3 experiments, with at least 500 cells counted per condition per experiment.

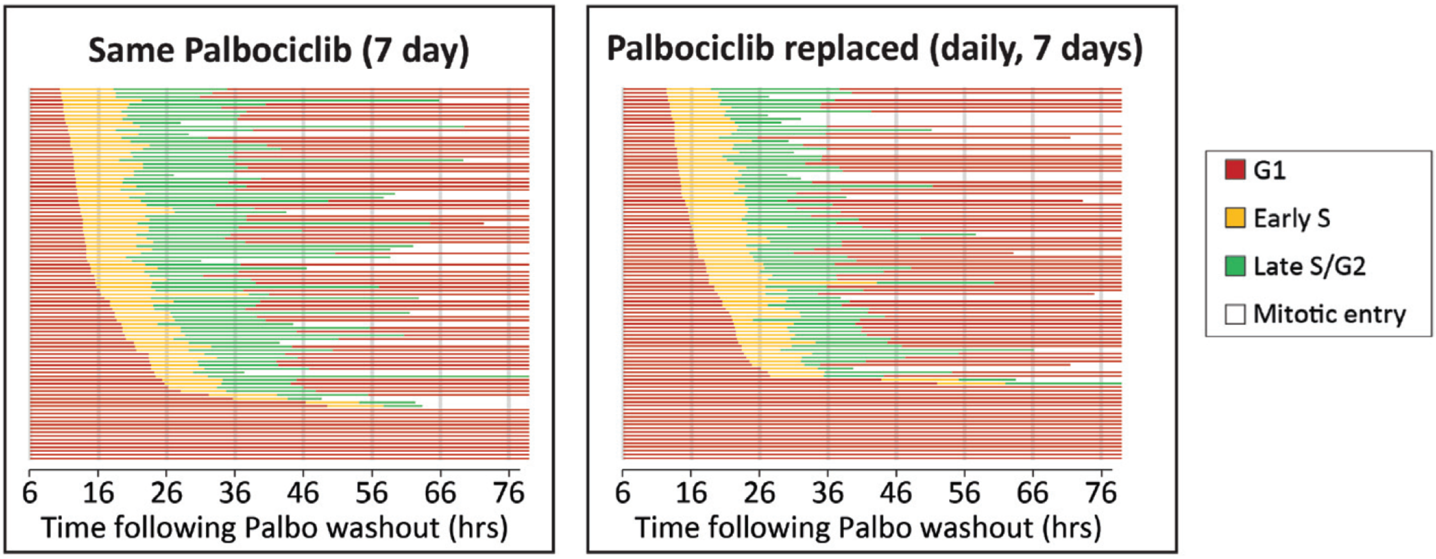


**Figure S3: No effect of replenishing the media on cell cycle profiles observed following a prolonged palbociclib arrest.** RPE1-FUCCI cells were treated with palbociclib (1.25 μM) for 7 days continuously (left panel) or refreshed daily with new media containing palbociclib (1.25 μM) for 7 days (right panel). Graphs show the cell cycle profile of individual RPE1-FUCCI cells (each bar represents one cell) after washout following the 7-day treatments. STLC (10 μM) was added to prevent progression through the first mitosis. 50 cells were analysed at random for each repeat and 3 experimental repeats are displayed (150 cells total).

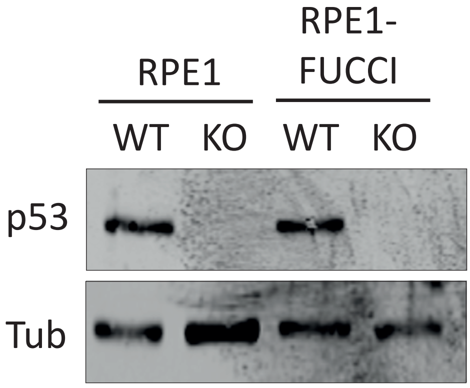


**Figure S4: Western to characterise the p53-WT and KO RPE1 and RPE1-FUCCI cells.** Western blot of whole cells lysates from p53-WT and KO RPE1 and RPE1-FUCCI cells treated with 50 μM Etoposide for 2 hours to induce p53.

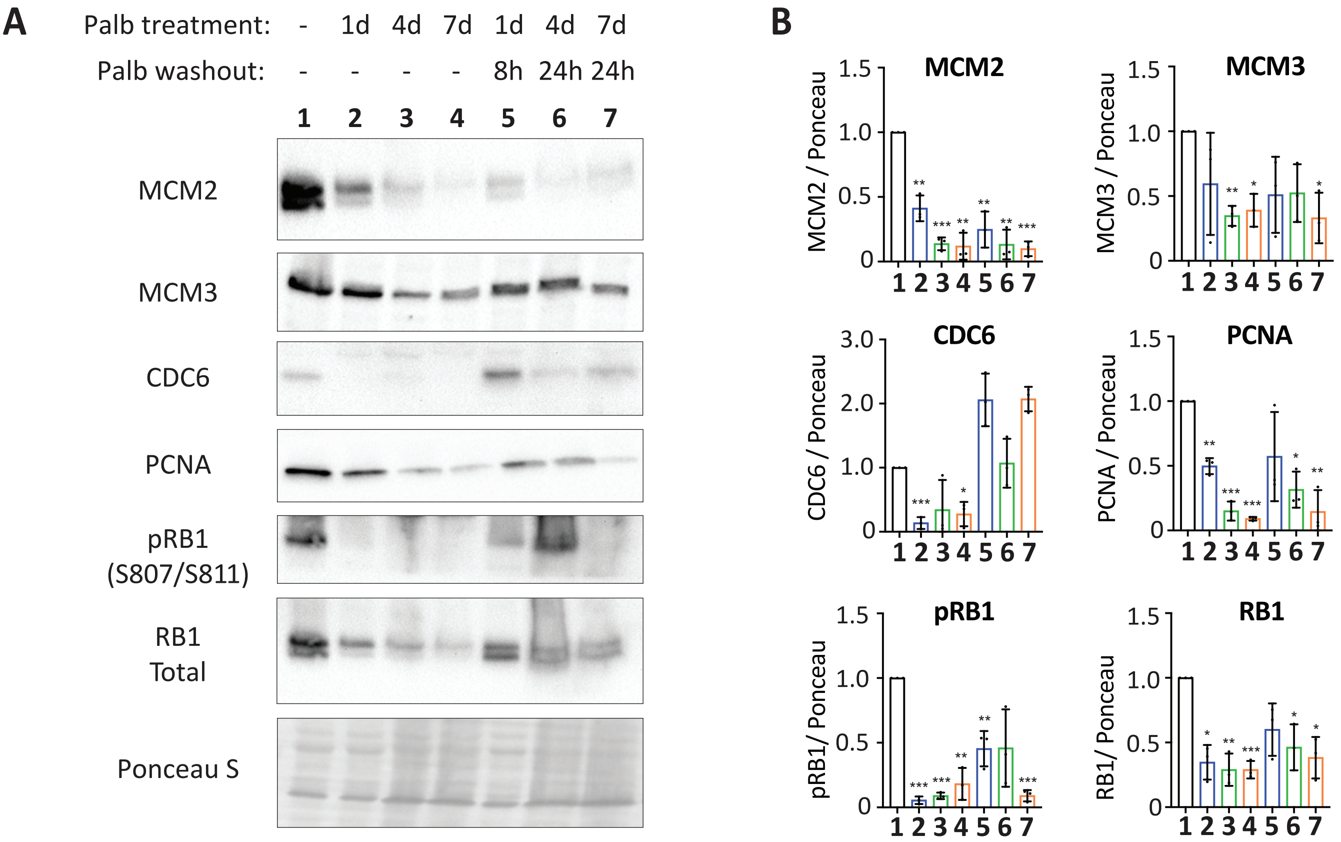


**Figure S5: Western blots to characterise the p53-WT and KO RPE1 and RPE1-FUCCI cells. (A)** Representative western blots of whole cell lysates from p53-KO RPE1 cells treated with palbociclib (1.25 μM) for 1, 4 or 7 days, or treated identically, and then washed out for the indicated times to reflect when the majority of cells are in S-phase (see Figure 1C). **(B)** Analysis of adjusted relative density from 3 independent western blot experiments. Bars display mean values -/+ SD. Significance determined by unpaired Student's T test comparing treated target protein to asynchronous target control. (* < 0.01, ** < 0.001, *** < 0.0001).

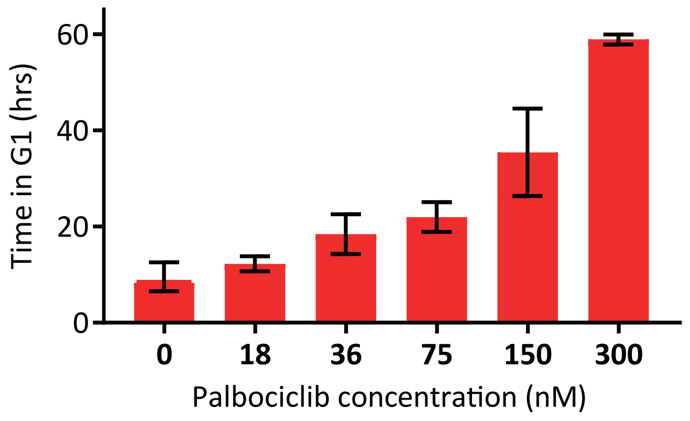


**Figure S6: Low doses of palbociclib progressively increase G1 length.** RPE1-FUCCI cells treated with low doses of Palbociclib (0-300nM) and the length of the first G1 after division was calculated based on the duration of mKO2-Cdt1 expression. Graphs display mean data from 2 experiments, with 50 cells analysed per condition per experiment.
